# Dynamic enhancer interactome promotes senescence and aging

**DOI:** 10.1101/2023.05.22.541769

**Authors:** Lu Wang, Gregory Donahue, Chen Zhang, Aaron Havas, Xue Lei, Caiyue Xu, Wenliang Wang, Golnaz Vahedi, Peter D. Adams, Shelley L. Berger

**Affiliations:** Penn Epigenetics Institute, Department of Cell and Developmental Biology, Perelman School of Medicine, University of Pennsylvania, Philadelphia, PA 19104, USA; Center for Translational Medicine, Thomas Jefferson University, Philadelphia, PA 19107, USA; Sanford Burnham Prebys Medical Discovery Institute, San Diego, CA, 92037, USA; Department of Genetics, University of Pennsylvania Perelman School of Medicine, Philadelphia, PA 19104, USA

## Abstract

Gene expression programs are regulated by enhancers which act in a context-specific manner, and can reside at great distances from their target genes. Extensive three-dimensional (3D) genome reorganization occurs in senescence, but how enhancer interactomes are reconfigured during this process is just beginning to be understood. Here we generated high-resolution contact maps of active enhancers and their target genes, assessed chromatin accessibility, and established one-dimensional maps of various histone modifications and transcription factors to comprehensively understand the regulation of enhancer configuration during senescence. Hyper-connected enhancer communities/cliques formed around genes that are highly expressed and within essential gene pathways in each cell state. In addition, motif analysis indicates the involvement of specific transcription factors in hyper-connected regulatory elements in each condition; importantly, MafK, a bZIP family transcription factor, was upregulated in senescence, and reduced expression of MafK ameliorated the senescence phenotypes. Because the accumulation of senescent cells is a key feature of aging, we further investigated enhancer connectomes in the liver of young and aged mice. Hyper-connected enhancer communities were identified during aging, which regulate essential genes that maintain cell differentiation and homeostasis. These findings reveal that hyper-connected enhancer communities correlate with high gene expression in senescence and aging and provide potential hotspots for therapeutic intervention in aging and age-associated diseases.

## Introduction

Cellular senescence is an irreversible cell-cycle arrest state induced by various stimuli, such as oncogene activation, DNA damage, and replication stress^1^. The accumulation of senescent cells during aging and in age-associated pathology, impairs tissue regeneration and repair capability in response to external stimuli^2^. This leads to the functional decline of tissues, and the clearance of senescent cells is critical for healthy aging. As cells become senesce, in addition to ceased proliferation, they acquire changes in genome organization, the chromatin landscape and gene expression^3-5^.

One important feature of senescence is the secretion of many proinflammatory factors, such as cytokines and chemokines, termed the senescence-associated secretory phenotype (SASP)^6^. Many SASP proteins promote inflammation and tissue damage, contributing to the development of age-associated disease. Approaches targeting SASP have significant therapeutic potential for age-related diseases^7^.

Gene expression programs are precisely regulated by the functional interactions between enhancers and their target genes, through the organization of the genome^8^. Global remodeling of the enhancer landscape occurs in senescence and aging^3^. Instead of influencing the nearest genes, many functional regulatory enhancer elements are located up to hundreds of kilobases away from their target genes^9^. In addition, dramatic genome reorganization occurs in senescent cells, such as forming senescence-associated heterochromatin foci (SAHF), and affecting the active regulatory interactions in euchromatin^10-13^. Therefore, it is critical to understand the influence of 3D regulatory change and enhancer interactions in senescence and how they are regulated.

Hyperconnected enhancer communities have been indicated to regulate essential genes in a context-dependent manner, in several biological systems^14-16^. These hyperconnected enhancer communities include multiple enhancers and target genes, and are thought to provide an important regulatory level in gene expression in steady state. Interestingly, cell identity genes are associated with and regulated by these hyperconnected enhancer communities^16^. Few studies investigate how these enhancer communities change in response to stress. As stress responsive gene programs and SASP gene activation are critical features of senescence and aging, there is a great need to understand the mechanisms underlying the dynamic alternations of these enhancer apparatus and how they contribute to senescence, aging and age-associated diseases.

Cellular stress during senescence and aging leads to altered gene expression and engagement of new enhancers, including upregulation of the SASP gene program^12, 13,17^. In this study, we mapped the enhancer connectome using H3K27ac HiChIP^18^, which captures enhancer- and promoter-centered contacts across the genome, and examined the dynamic interplay between 3D enhancer networks and gene transcription in senescence induced by oncogene activation. We find that the hyperconnected enhancer communities are dynamic in senescence. The alteration of these enhancer communities in senescent cells is mediated by MafK, a bZIP family transcription factor. Knockdown of MafK can ameliorate senescence phenotypes, including reduced β-galactosidase (β-gal) accumulation and lower SASP gene expression. Moreover, cut&run profiles confirm the binding of MafK at SASP genes increases in senescent cells. Furthermore, we identify hyperconnected enhancer communities in hepatocytes isolated from young and aged mice, which encompass genes crucial for age-specific physiological states. 3D enhancer landscape in aged hepatocytes is wired to a disease-susceptible state. Together, our data indicate that hyperconnected enhancer communities drive transcriptional changes during senescence and aging.

## Results

### Chromatin accessibility is altered in senescence

We utilized primary human lung fibroblasts (IMR90) and induced senescence of these cells by expression of oncogenic HRasV12 (Supp Fig 1A). Senescence is associated with dramatic changes in genome organization and epigenetic landscape^3, 19^. To assess chromatin changes, we first measured the chromatin accessibility using ATAC-seq (assay for transposase-accessible chromatin with high-throughput sequencing)^20^ in both proliferating and senescent cells (Supp Fig 1A). Biological replicates show great reproducibility (Supp Fig 1B). Approximately 128K open chromatin regions were identified, among which ∼17K regions significantly increased accessibility in senescence, while ∼12K significantly reduced accessibility in senescence (Fig 1A).

**Figure 1.**
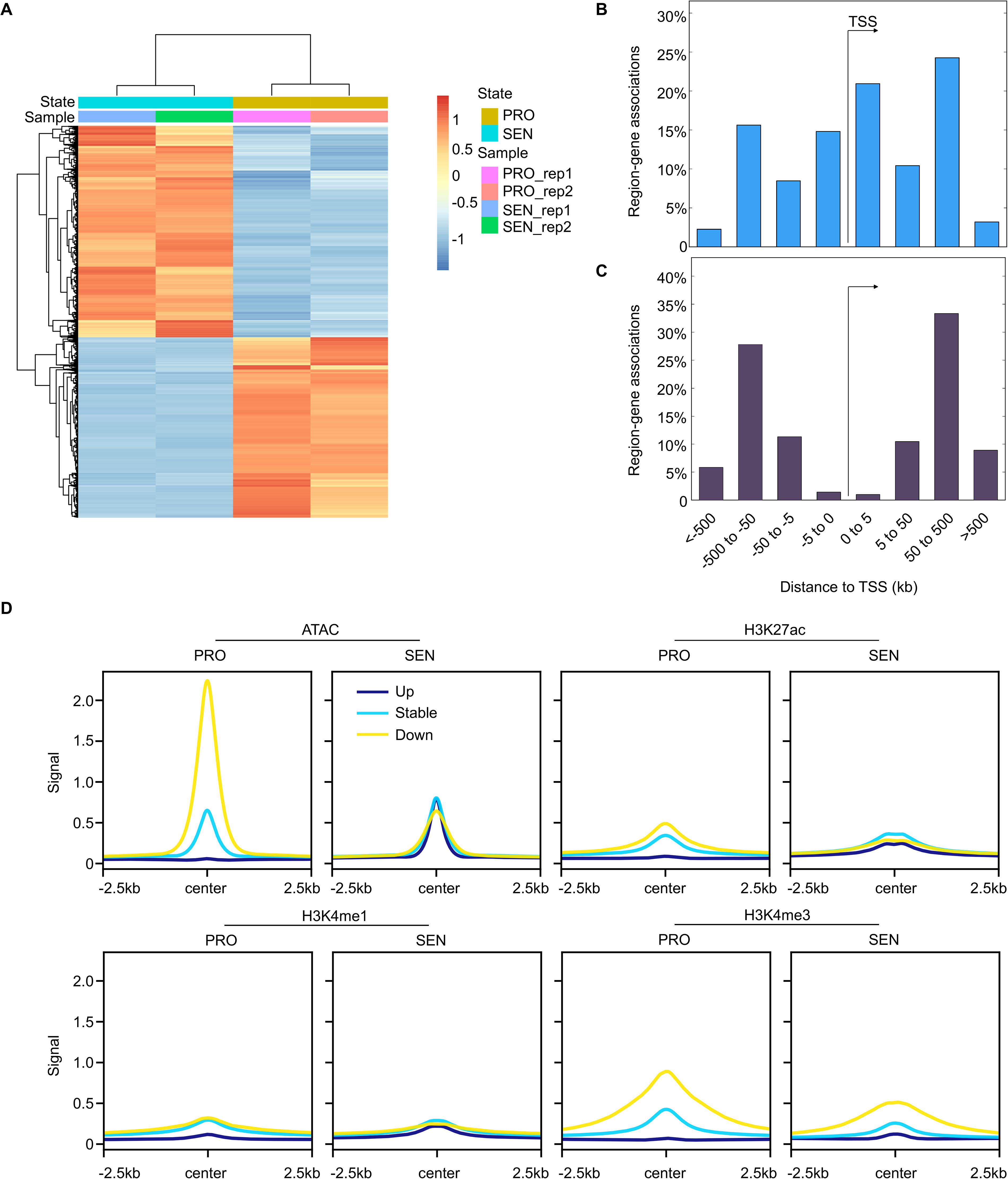
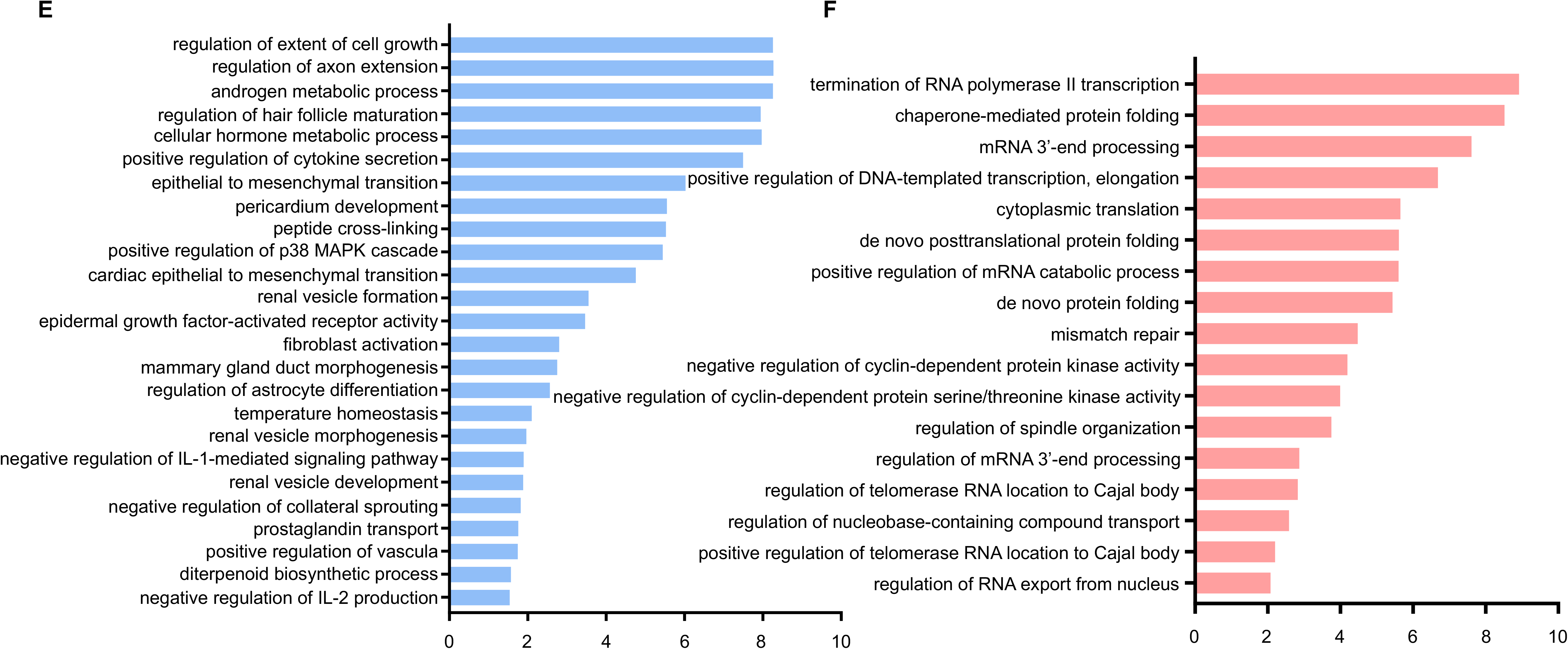
Chromatin accessibility is altered in senescence. A. Heatmaps depicts the differential ATAC-seq peaks in proliferating and senescent cells. B. Bar-plot demonstrates the distribution of the distance of differentially downregulated ATAC-seq peaks in senescence. C. Bar-plot demonstrates the distribution of the distance of differentially upregulated ATAC-seq peaks in senescence. D. Metaplots demonstrate the changes of histone modifications – H3K27ac, H3K4me1 and H3K4me3 in up-, down-regulated and stable ATAC-seq peaks in senescence. E. Gene ontology terms enriched in down-related ATAC-seq peaks in senescence from GREAT analysis. F. Gene ontology terms enriched in up-regulated ATAC-seq peaks in senescence from GREAT analysis. PRO, proliferating cells; SEN, senescent cells; rep, replicate; TSS, transcription start site.

To further understand the biological significance of these alterations in chromatin accessibility, we examined the distance of these open chromatin regions to gene transcription start sites (TSSs). Regions with reduced accessibility were located mainly within 50 kb up- and down-stream of TSSs (Fig 1B), while most regions with increased accessibility in senescence are located at distal regulatory regions (50-500 kb) to TSSs (Fig 1C). These findings suggest that in the progression from proliferation to senescence, distal regulatory elements, including enhancers, may play critical roles in activating senescence-specific gene programs in response to external stimuli. Enhancers and promoters possess distinct features of histone modifications. Mono-methylation of histone 3 at lysine 4 (H3K4me1) and acetylation of histone 3 at lysine 27 (H3K27ac) are preferentially enriched at active enhancers, while tri-methylation of histone 3 at lysine 4 (H3K4me3) is mostly enriched at promoters^21^. Indeed, the distal regulatory regions that increased accessibility in senescence gained H3K27ac and H3K4me1 (Fig 1D, upper panel), suggesting these regions are mainly enhancers. The regions with reduced accessibility in senescence were mostly marked by H3K4me3 (Fig 1D, lower panel), suggesting a promoter identity.

To gain functional insights of these alterations in chromatin accessibility, we performed Gene Ontology (GO) analysis. Genes regulated by regions with increased accessibility in senescence are enriched in positive regulation of cytokine secretion and p38 MAPK cascade (Fig 1E), indicating that these enhancer regions regulate the senescence-specific pathways. Genes regulated by regions with reduced accessibility in senescence are enriched in transcription regulation and cyclin-dependent protein kinase activity (Fig 1F), suggesting a role in cell proliferation.

### Distinct regulatory programs are mediated by different transcription factors in proliferating and senescent cells

Active enhancers are transcribed yielding enhancer RNAs (eRNAs). ATAC-seq data unveiled the global changes of chromatin accessibility in senescence, but the information on the transcriptional activity of these changing regions is not provided. To further understand the function of these elements, we assessed the activity of these regulatory elements. Precision nuclear run-on sequencing (PRO-seq) detects nascent RNAs with high sensitivity^22^. We performed PRO-seq in both proliferating and senescent cells. Biological replicates show good reproducibility (Supp Fig 1C-D).

PRO-seq data also provides insights into gene expression, and thus we analyzed transcription within annotated protein-coding genes and regulatory elements^23^. In line with previous reports using RNA-seq^12, 24^, SASP genes were significantly upregulated in senescence. In total, in PRO-seq data, 714 genes were upregulated in senescence, and 533 were downregulated (Fig 2A). In addition, active transcriptional regulatory elements (TREs), including promoters and enhancers, were identified in both conditions. To further identify transcription factors (TFs) that may be involved in maintaining each regulatory program (proliferation or senescence), we identified enriched TF motifs in differentially regulated TREs. TF motifs enriched in down-regulated TREs were bZIP-ATF, E2F, and zinc finger-containing transcription factors (Fig 2B). TF motifs enriched in up-regulated TREs were ETS family and nuclear factor Kappa B (NF-kB) (Fig 2C). These identified TFs were mutually exclusive, suggesting that different groups of TFs mediate distinct regulatory programs in proliferating and senescent cells.

**Figure 2.**
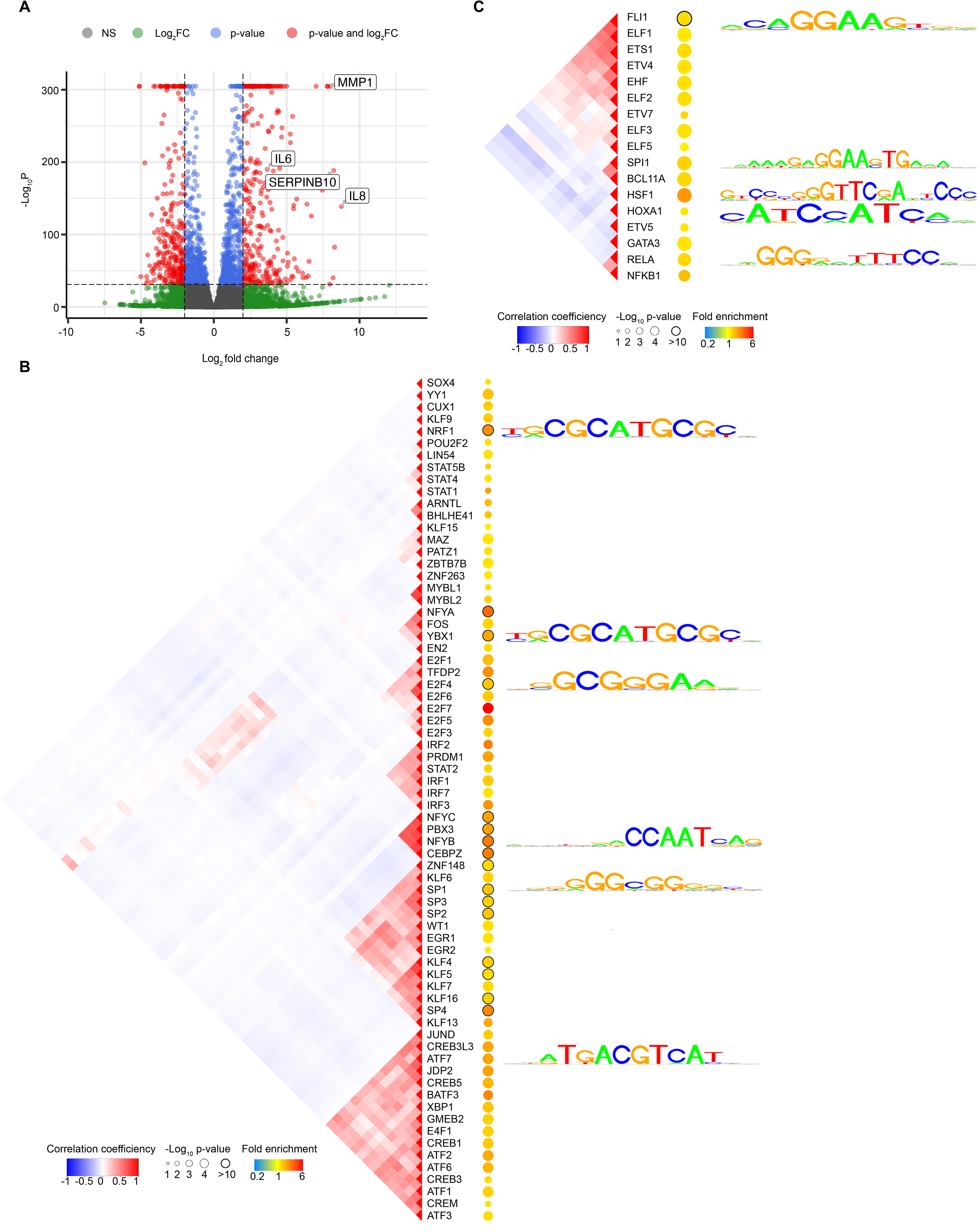
Nascent transcription alters in senescence. A. Volcano plot demonstrates the differentially expressed genes in PRO-seq data. B. Transcription factors enriched in down-regulated transcriptional regulatory elements in senescence. C. Transcription factors enriched in up-regulated transcriptional regulatory elements in senescence.

### Enhancer interactome influences gene transcription

Functional enhancers can be located hundreds of kilobases away from their target genes^25^. Given the significance of enhancers in senescence, we investigated changes in the enhancer interactome and their influence on gene expression. We performed H3K27ac HiChIP to identify the enhancer interactome in proliferating and senescent cells (Supp Fig 1A). H3K27ac HiChIP is a highly efficient and sensitive method to capture enhancer- and promoter-centered contacts across the genome^18^. We identified ∼240K and 200K interactions in each condition (Supp Fig 2A-B). The median interaction distance between anchoring regions ranged from 220 to 225 kb (Supp Fig 2A-B). Most of the interactions are enhancer-enhancer contacts (Supp Fig 2C). We found multiple potential enhancers can regulate one single gene. We found that the average interactions per promoter was 4 to 5 (Fig 3A-B) in both proliferating and senescent cells.

**Figure 3.**
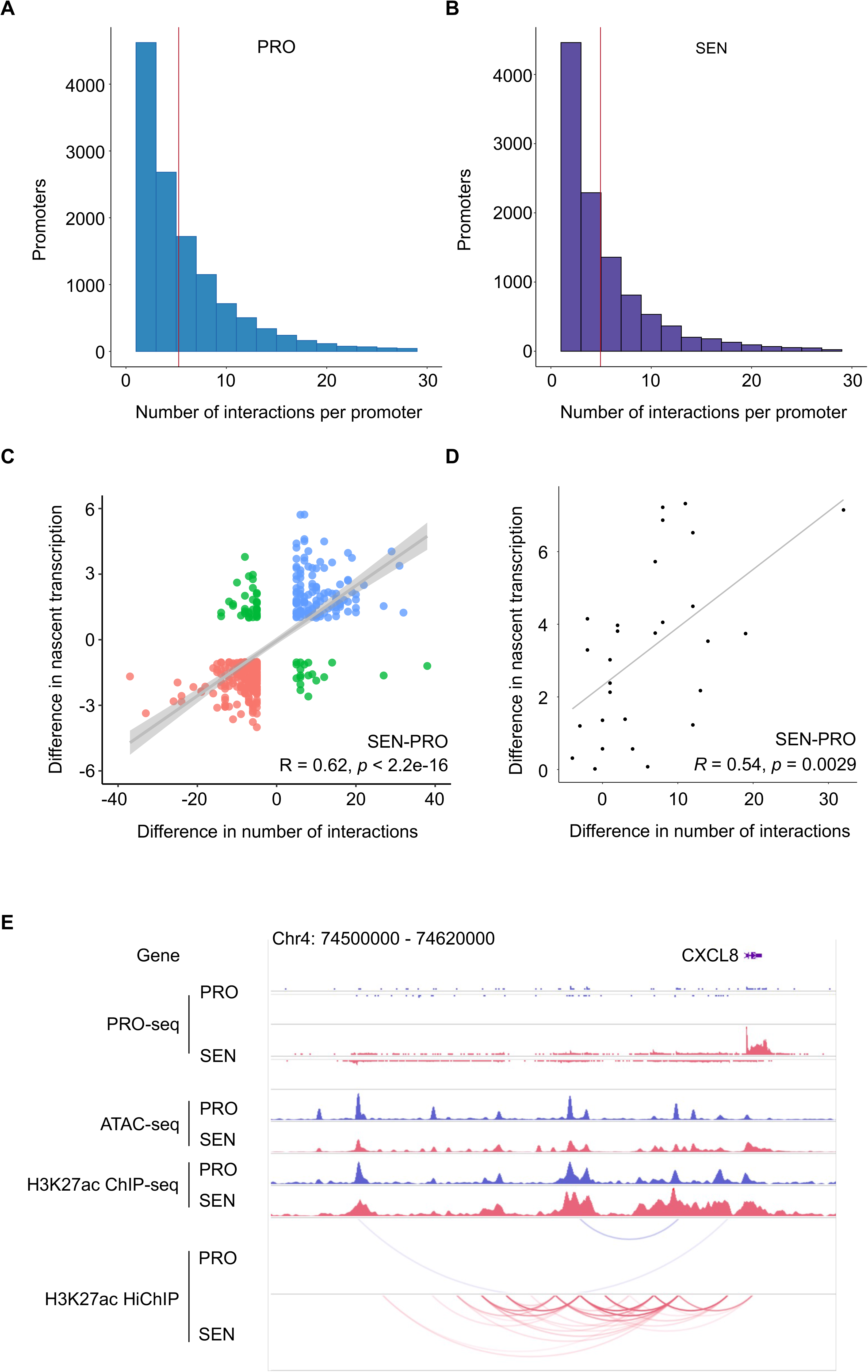
Multiple regulatory elements regulate gene expression. A. Bar-plot demonstrates the number of interactions contacting each promoter in proliferating cells. Green, genes with higher expression level in proliferating cells; Red, genes with higher expression level in senescent cells. B. Bar-plot demonstrates the number of interactions contacting each promoter in senescent cells. Green, genes with higher expression level in proliferating cells; Red, genes with higher expression level in senescent cells. C. Scatter plot depicts the correlation between the difference in the number of interactions and the difference in nascent transcription. Pearson correlation coefficient is calculated. D. Scatter plot depicts the correlation between the difference in the number of interactions and the difference in nascent transcription of senescence-associated secretory phenotype (SASP) genes. Pearson correlation coefficient is calculated. E. The genome browser view demonstrates the nascent transcription (PRO-seq), chromatin accessibility (ATAC-seq), H3K27ac ChIP-seq, as well as H3K27ac HiChIP interactions at the CXCL8 locus. PRO, proliferating cells; SEN, senescent cells.

Enhancer regulation can control gene expression temporally or spatially in response to environmental cues^26^. To connect global chromatin alterations with gene transcription in senescence, we compared nascent transcription with the number of regulatory elements interacting with each promoter. The alterations in the number of chromatin interactions contacting gene promoters corresponded positively with gene transcription changes (Fig 3C), suggesting that 3D enhancer regulation may have a key role in regulating the cellular and molecular response to stress. Specifically, most of SASP genes were regulated by multiple regulatory elements via gaining interactions in senescence, and the number of gained interactions positively correlated with their expression (Fig 3D). For example, the pro-inflammatory cytokine IL8 is an important component of SASP, and multiple enhancers specifically contacted the promoter of IL8 in senescent cells, which corresponds to the increased IL8 transcription (Fig 3E).

### State-specific genes with hyper-connected enhancer communities

Multiple regulatory elements control gene expression, and recent studies have shown that hyper-connected enhancer communities control important lineage- and disease-specific genes^14-16^. Therefore, we set out to understand the role of these hyper-connected enhancer communities in senescence. We identified a set of promoters with increased levels of chromatin interactivity. Our analyses showed that hyper-connected enhancer communities form in both proliferating and senescent cells, and state-specific genes are regulated by these hyper-connected enhancer communities (Fig 4A-B). For example, key state-specific genes regulated in this way include COL1A2 and OSR1 for proliferating fibroblasts, and HMGA2, SERPINB2 and CXCL2 for senescence (Fig 4A-B), and their expression correlates with presence in specific hyper-connected enhancer communities. Specifically, in proliferating cells, genes regulated by these communities are enriched in proliferation and fibroblast function-related pathways, such as regulation of proliferation (Fig 4C); in senescent cells, genes regulated by these communities are enriched in senescence-responsive pathways, such as regulation of inflammatory response and ERK1 and ERK2 cascade (Fig 4D). These pathways are activated in senescence, and play an essential role in establishing and maintaining the senescence phenotypes^6, 27^.

**Figure 4.**
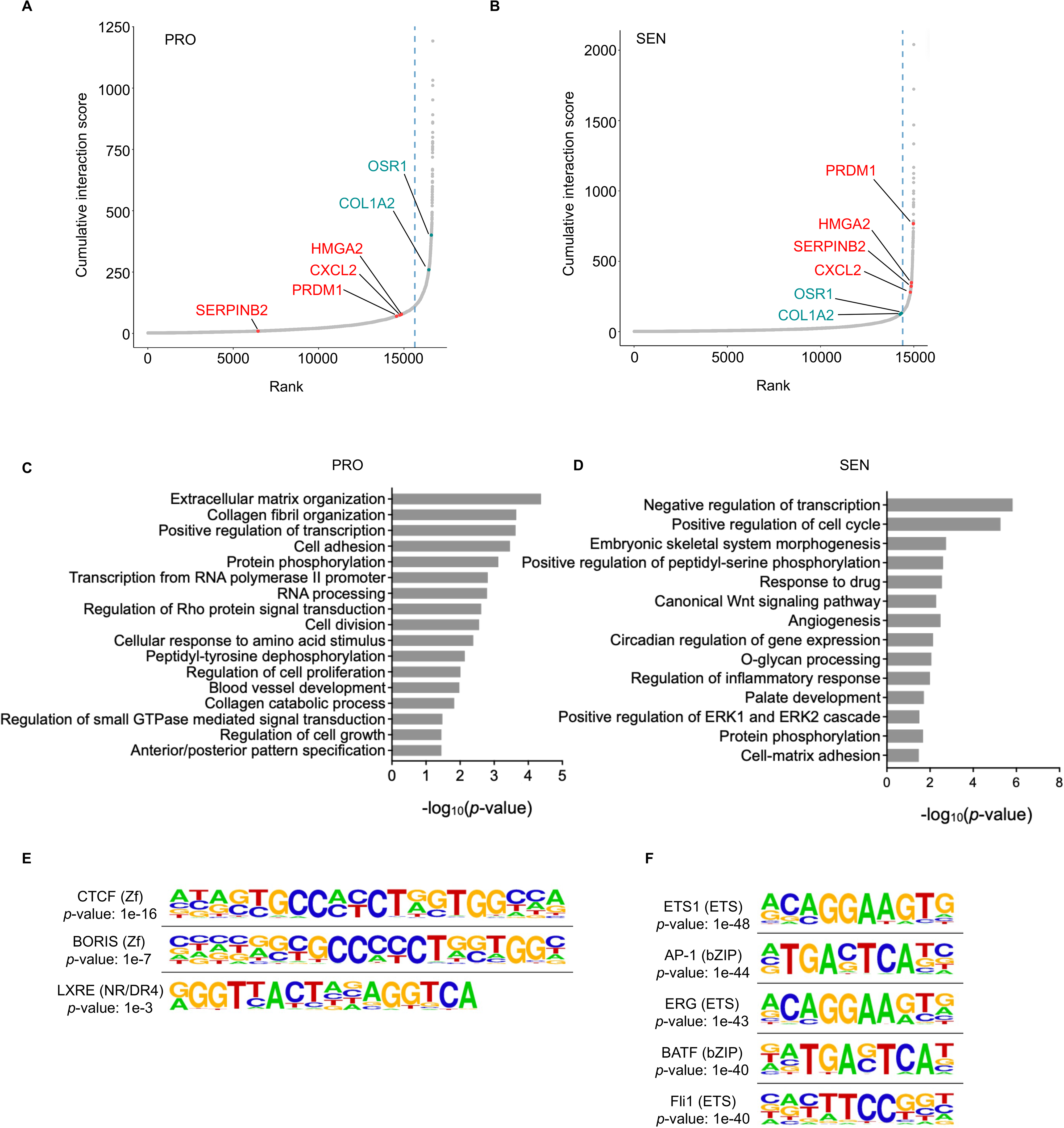
Hyper-connected enhancer communities regulate state-specific genes. A. Plot depicts the number and strength of the interactions in hyper-connected enhancer communities in proliferating cells. B. Plot depicts the number and strength of the interactions in hyper-connected enhancer communities in senescent cells. C. Gene ontology terms enriched for genes regulated by proliferation-specific enhancer communities. D. Gene ontology terms enriched for genes regulated by senescence-specific enhancer communities. E. Motifs enriched in proliferation-specific enhancer communities. F. Motifs enriched in senescence-specific enhancer communities. PRO, proliferating cells; SEN, senescent cells.

To gain insights into the underlying mechanisms of the formation of these 3D enhancer communities, we performed motif analysis of DNA sequences within regulatory elements involved specifically in each state. The most enriched motif in proliferating cells was CTCF (Fig 4E), consistent with its universal function in mediating enhancer-promoter interactions. However, CTCF was not enriched in senescence-specific enhancer communities (Fig 4F). Indeed, as previously observed^28^, CTCF protein level was downregulated in senescent cells, yet most of the CTCF binding sites maintain in senescent cells (Supp Fig 2D). This suggests that in senescent cells, in addition to common structural proteins, other transcription factors, including ETS and AP-1 families, may significantly contribute to the connectivity of the enhancer communities to regulate senescence-specific gene programs.

### AP-1 transcription factors in regulating hyper-connected enhancer communities

Senescent cells accumulate in aged tissues contributing to age-associated diseases, and the clearance of senescent cells promotes healthy aging^29^. Therefore, we focused on the transcription factors that are specifically enriched in senescence. The Activator Protein 1 (AP-1) motif was among the top enriched TFs (Fig 4F), and it has been reported to play critical roles in senescence across conditions^30, 31^. It is involved in stress response and many other physiological conditions. Further, AP-1 is a key regulator of enhancer function. The AP-1 family is composed of different subunits, including FOS, JUN, MAF (musculoaponeurotic fibrosarcoma) and ATF (activating transcription factor) protein families, and they share the core motif sequence^32^. These family members can form homo- and hetero-dimers to exert different functions.

To understand the role of AP-1 in regulating the hyper-connected enhancer communities in senescence, we first examined the protein level of different AP-1 family members in proliferating and senescent cells; most were upregulated in senescent cells (Fig 5A). Small MAF proteins have been implicated in stress response, inflammation and disease^33, 34^. We carried out MafK Cut&Run, a sensitive assay for mapping protein-DNA interactions^35^. We found that MafK binding sites increased in hyper-connected enhancer communities in senescent cells (Fig 5B-C), which is positively correlated with gene expression changes. Interestingly, we found that MafK binding sites significantly increased in hyper-connected enhancer communities that regulate SASP genes (Fig 5D). For example, MafK binding sites increased around the IL1B gene, a key SASP gene, and these sites form a senescence-specific hyper-connected enhancer community (Fig 5E). This suggests MafK may be involved in regulating senescence phenotypes through these hyper-connected enhancer communities.

**Figure 5.**
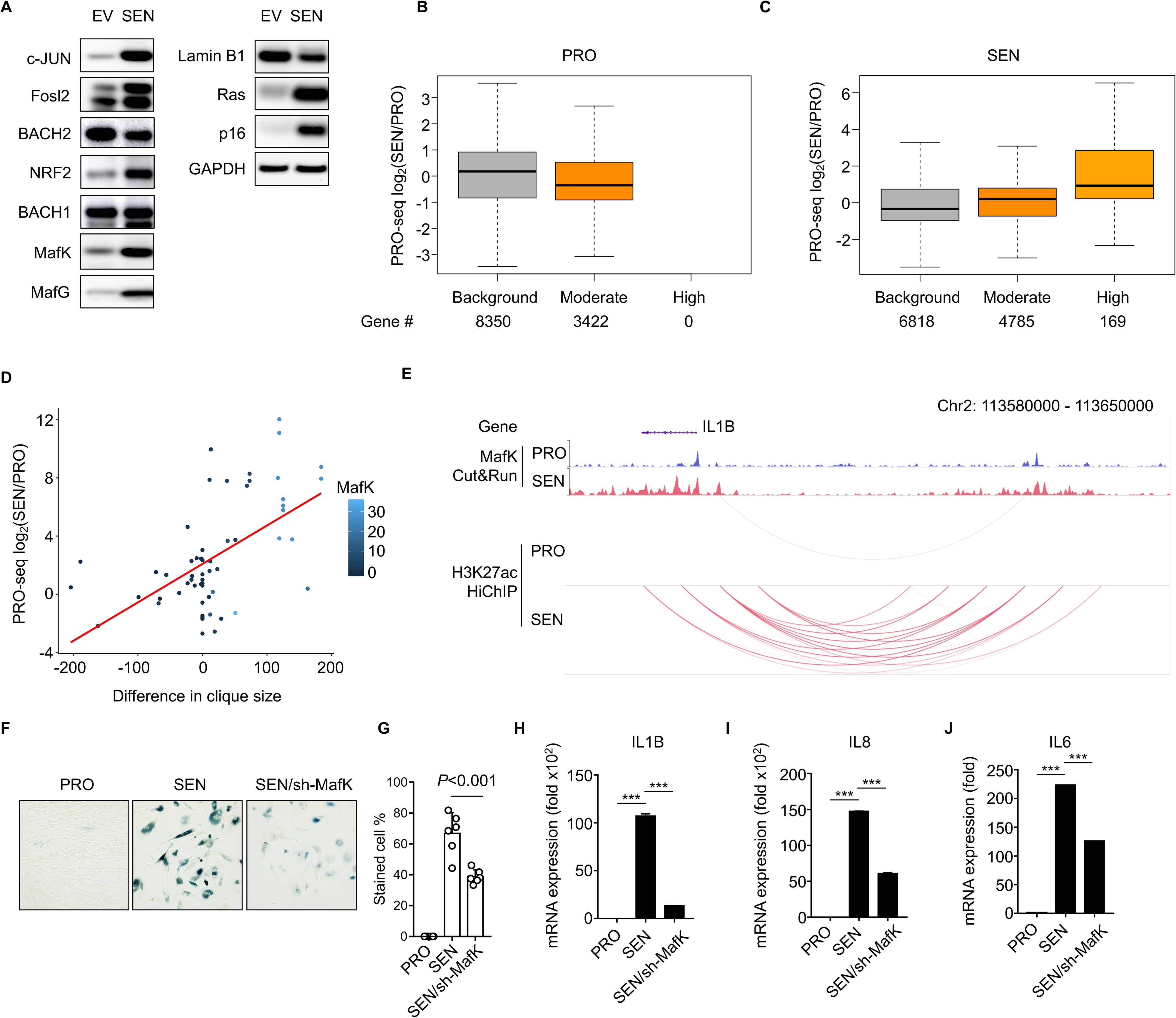
MafK regulates enhancer communities and senescence phenotypes. A. Protein levels of bZIP family transcription factors in control and senescent cells. EV, empty vector; SEN, senescent cells. Lamin B1, Ras, and p16 serve as controls to show the cells are senesced. B. Boxplot demonstrates that differentially expressed genes classified according to their number of pro-specific loops with pro-specific MafK peaks show no relationship between MafK binding and gene expression. C. Boxplot demonstrates that differentially expressed genes classified according to their number of SEN-specific loops with SEN-specific MafK peaks show a regulatory bias, where more newly looped MafK peaks produce higher upregulation in SEN. D. Scatter plot depicts changes in MafK binding in enhancer communities regulating SASP genes, suggesting the highly upregulated SASP genes with increased enhancer contacts are influenced by many OIS-specific MafK binding events. E. The genome browser view of MafK binding sites (Cut&Run), and enhancer interactions (H3K27ac HiChIP) around the IL1B locus. F. Beta-gal staining results in proliferating, senescent and senescent with MafK knockdown cells. It shows reduced MafK level leads to alleviated senescence phenotype. G. Quantifications for the beta-gal staining experiments in F. SASP gene expression, IL1B (H), IL8 (I), IL6 (J), quantified by RT-qPCR in proliferating, senescent, and senescent with MafK knockdown cells. PRO, proliferating cells; SEN, senescent cells; sh-MafK, knockdown of MafK by shRNAs.

To further test whether MafK has function in senescence, as proof of concept, we knocked down MafK by shRNA and examined senescence phenotypes. We found that when MafK protein level is reduced, the senescence phenotype was alleviated by beta-galactosidase staining (Fig 5F), with quantification in Fig 5G. Furthermore, knockdown of MafK could decrease the expression of several SASP genes, which are regulated by senescence-specific hyper-connected enhancer communities (Fig 5 H-J). Together, these results indicate MafK influences senescence-responsive genes potentially through regulating hyper-connected enhancer communities.

### Disease-associated traits regulated by hyper-connected enhancer communities in senescence

Distal regulatory regions, such as enhancers, can harbor genetic variants associated with human complex traits identified by genome-wide association studies (GWASs)^36, 37^. As most of these genetic variants are located in non-coding regions, it is difficult to clearly identify their functional roles^36, 37^. Our high-resolution contact maps in senescence and aging can be investigated for genetic variations associated with aging and age-related diseases. As senescent cells contribute to the development of many age-related diseases, we interrogated whether specific traits are enriched in the senescence-specific enhancer communities. We found that several age-related diseases were more enriched in senescence-specific enhancer communities, such as type 2 diabetes, and pro-inflammatory response, such as high IL-1 beta levels in gingival crevicular fluid (Fig 6A).

**Figure 6.**
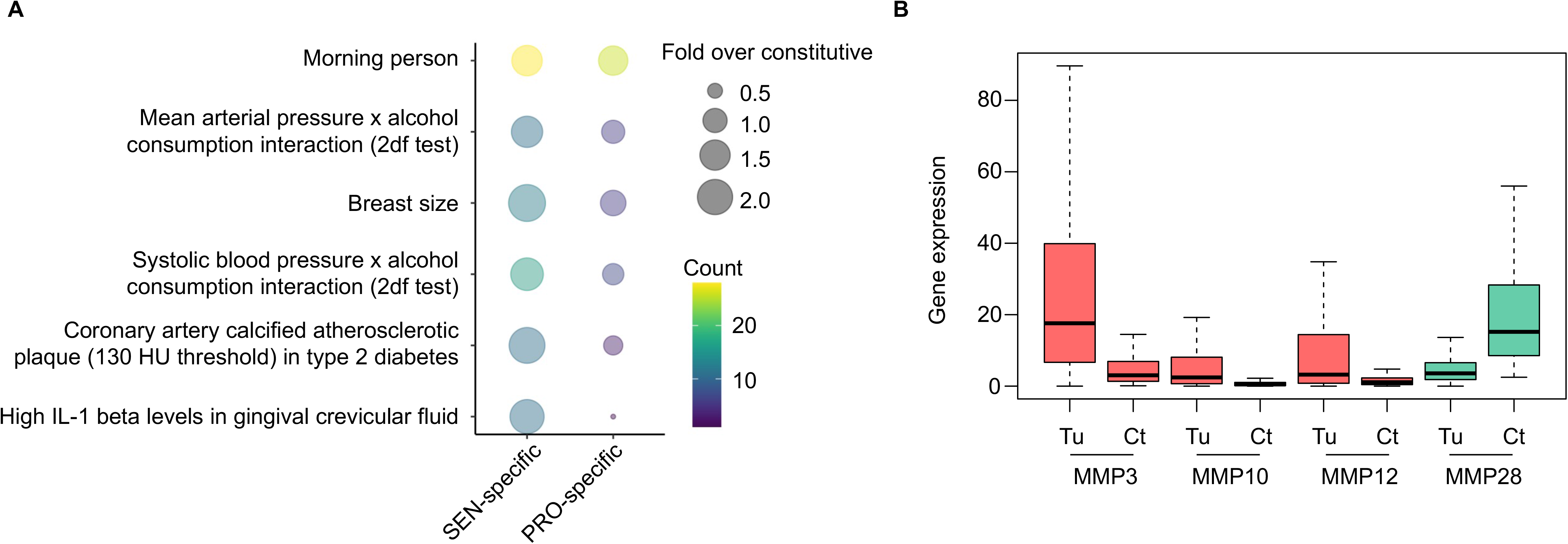
Disease-associated traits in hyper-connected enhancer communities. A. Bubble plot demonstrates GWAS traits associated with senescence-specific enhancer communities. GWAS terms enriched for SNP overlaps in SEN-specific loops, sorted from top-to-bottom by differential enrichment in SEN. B. Boxplot depicts expression levels of MMP genes, which are regulated by enhancer communities in senescence, in breast cancer patients and controls. Tu, tumor; Ct, control. SEN, senescent cells.

Aging is a primary risk factor for many diseases, including cancer. To explore whether hyper-connected enhancer communities contribute to the risk of cancer, we compared the data from the cancer genome atlas (TCGA) with our senescence-specific enhancer maps^38^. Matrix metalloproteinase (MMP) genes are involved in remodeling extracellular matrix and have been implicated in senescence and tumor invasion^39, 40^. Among the six MMP genes classified as SASP components, we found that MMP3, MMP10 and MMP12 showed increased interactions and expression in senescence and had higher expression level in breast cancer patients comparing to the control group (Fig 6B, Supp Fig 2E). Particularly, MMP3 has been linked to promoting breast cancer invasion^41, 42^. Notably, the non-SASP MMP gene, MMP28, which was not involved in enhancer communities, did not increase its expression in cancer samples (Fig 6B). These findings suggest that changes in enhancer communities in senescence affect key gene expression, which may further contribute to the development of breast cancer.

### Hyper-connected enhancer communities regulate genes essential to maintain rejuvenation in liver

To gain insights into organismal aging, we next investigated hyper-connected enhancer communities in hepatocytes isolated from young (age 5 months) and old mice (age 25 months). We performed H3K27ac HiChIP and identified hyper-connected enhancer communities in young and old hepatocytes (Supp Fig 3A-B). As with the proliferating and senescent cells, we found that age-specific enhancer communities regulate genes enriched in essential pathways reflecting the physiological state (Fig 7A-B). Specifically, genes that are critical for liver functions, such as L-serine metabolic process and fatty acid beta-oxidation^43, 44^, are associated with enhancer communities specifically in young mice (Fig 7A). In old mice, inflammatory response and cell chemotaxis pathways are specifically enriched (Fig 7B), consistent with the changes observed in liver during aging^45^. For example, cytochrome genes, important for metabolic functions in liver^46^, lose interactions (Fig 7C) and have reduced gene expression (Fig 7D) in old mice. Moreover, genes that are repressed in liver fibrosis^47^, were decreased in their expression in old hepatocytes (Fig 7E), and were associated with the loss of interactions (Fig 7F). This suggests that connected enhancers play essential roles in maintaining liver function and may be lost during aging and in age-related diseases. Similarly, we observed enhancer communities formed around disease-associated genes in aged hepatocytes, including CCND1 (Fig 7G).

**Figure 7.**
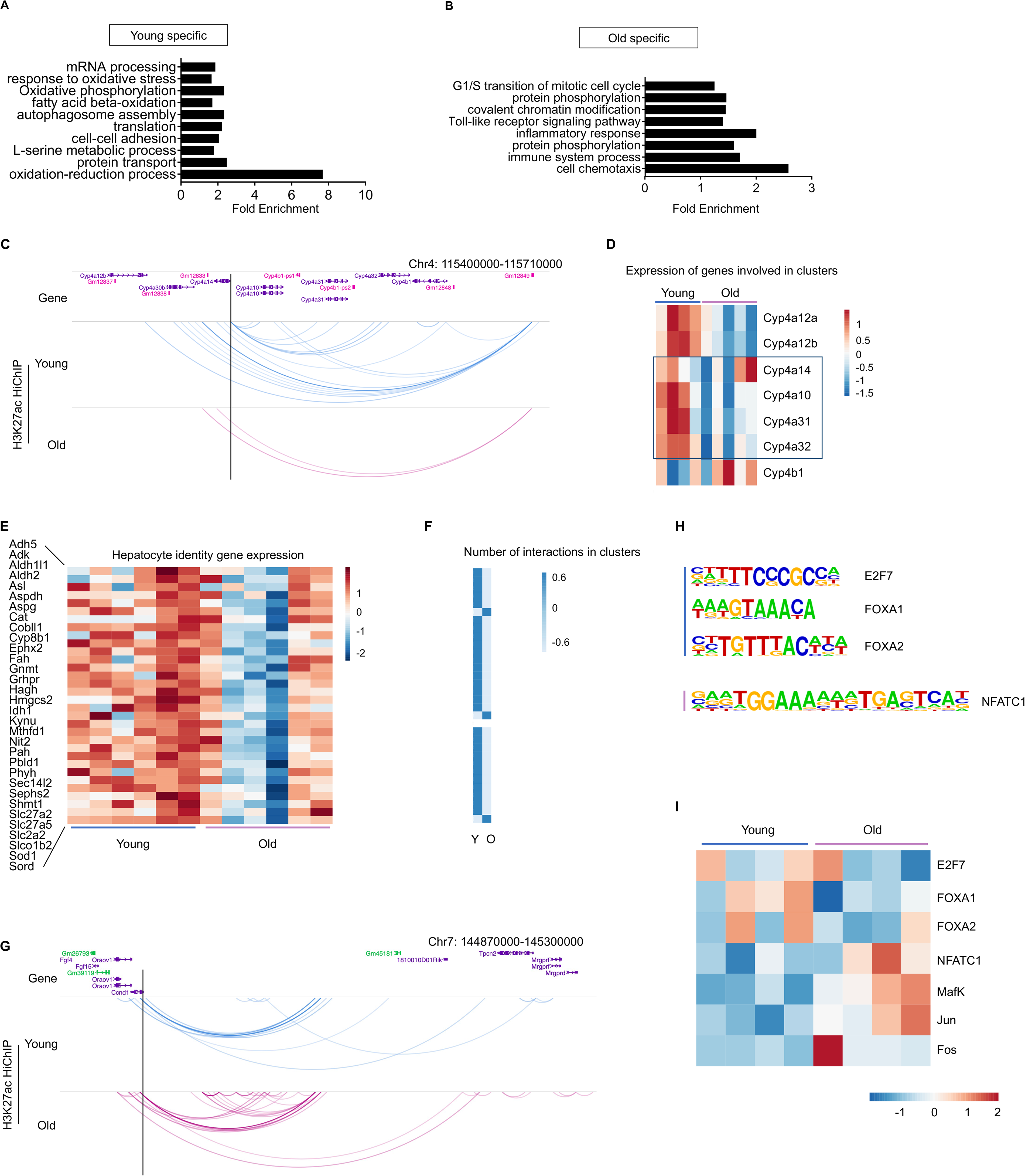
Hyper-connected enhancer communities are dynamic during aging. A. Gene ontology terms enriched for genes regulated by hyper-connected enhancer communities in hepatocytes from young mice. B. Gene ontology terms enriched for genes regulated by hyper-connected enhancer communities in hepatocytes from olde mice. C. The genome browser view of enhancer interactions around cytochrome genes in young and old mice. D. Heatmap demonstrates the changes of cytochrome genes involved in the loci showed in C. E. Heatmap depicts the expression levels of genes downregulated in fatty liver diseases, in young and old mice. F. Heatmap illustrates changes in enhancer interactions around the genes in E. G. The genome browser view of enhancer interactions around CCND1 locus in young and old mice. H. Motifs enriched in young-specific and old-specific hyper-connected enhancer communities. I. Heatmap demonstrates the expression levels of transcription factors regulating hyper-connected enhancer communities in young and old mice.

Motif analyses showed that distinct transcription factors were enriched in enhancer communities in young or old hepatocytes (Fig 7H), and they were relatively highly expressed in corresponding state (Fig 7I). Specifically, FOXA1 and FOXA2 were enriched in enhancer communities in young hepatocytes, and they are pioneer factors that can regulate enhancers and transcription^48^. In aged hepatocytes, NFATC1 was enriched, which belongs to bZIP family, and it has been indicated in promoting liver cell damage and inflammation^49^. Taken together, these data indicate that hyper-connected enhancer communities may regulate age-dysregulated genes, which may in turn be disease-priming.

## Discussion

Here, we set out to understand the mechanisms regulating enhancer interactions in senescence and aging. We generated high-resolution contact maps centering on enhancers and identified multi-enhancer interactions within communities controlling gene expression in a state-specific manner. Further we found that MafK, a bZIP family transcription factor, mediates the alteration of these enhancer communities in senescent cells, which further influence SASP gene expression and senescence phenotypes. Additionally, we found that these enhancer interactome alterations in senescence are associated with age-related disease traits, such as such as type 2 diabetes and high IL1 beta levels in gingival crevicular fluid. We also found that several matrix metalloproteinase genes have altered enhancer interactions in senescence which are associated with their upregulation in breast cancer. These findings emphasize the potential role of senescence in disease development through understanding their 3D enhancer connectomes, and potentially provide new therapeutic targets to prevent age-related diseases.

Recent studies showed that hyperconnected enhancer communities regulate essential genes in a context-dependent manner^14-16, 50^. In addition to steady state and selected loci, we identified genome-wide dynamic hyperconnected enhancer communities in proliferating and senescent cells, hepatocytes isolated from young and old mice. We found that genes that are essential to maintain liver functions lost interactions in old mice, and the enhancer connectome in old mice has already acquired signatures of age-related diseases, such as liver fibrosis and liver cancer. These results suggest that enhancer communities change in response to stress, and these alterations contribute to the phenotypes in the new state, which could serve as a new important layer of regulation and a pathway to target for therapeutics.

In sum, our results emphasize the importance of understanding the dynamic interplay between 3D enhancer networks and gene transcription in senescence and aging, as this knowledge may lead to the development of novel interventions to delay or prevent age-related diseases. Our work suggests that targeting hyperconnected enhancer communities and MafK-mediated enhancer rewiring may provide new therapeutic strategies to treat age-related diseases. Future studies are necessary to further explore this exciting direction.

## Figure legends

**Supplemental Figure 1.**
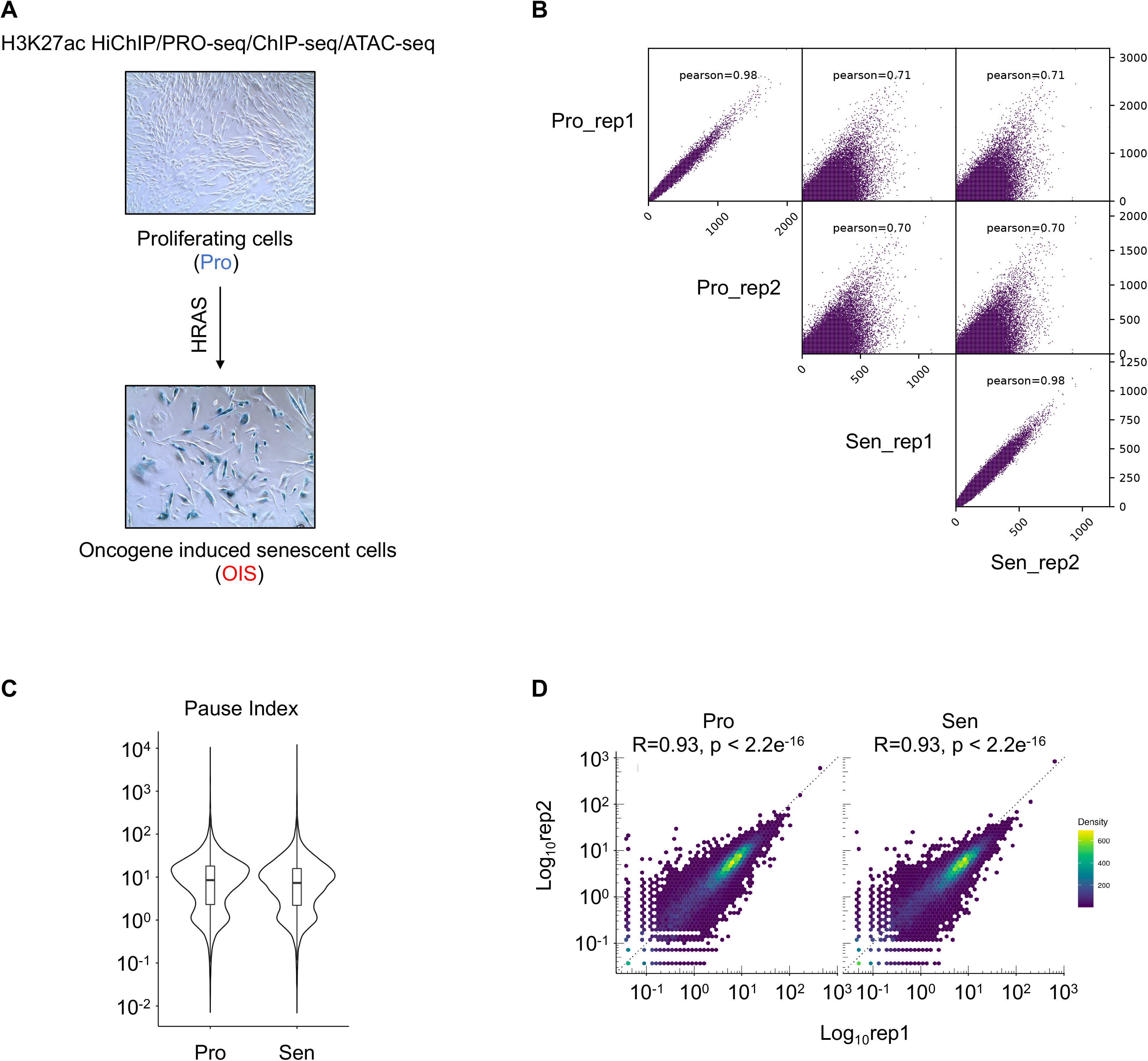
Overview of the study and data assessment. A. Scheme of the experiments conducted in this study. B. Reproducibility of ATAC-seq libraries. C. Distribution of pause index in qPRO-seq samples. D. Correlation of reads-per-million (RPM) in promoter regions between replicates. Pro, proliferating cells; Sen, senescent cells; rep1, replicate 1; rep2, replicate 2.

**Supplemental Figure 2.**
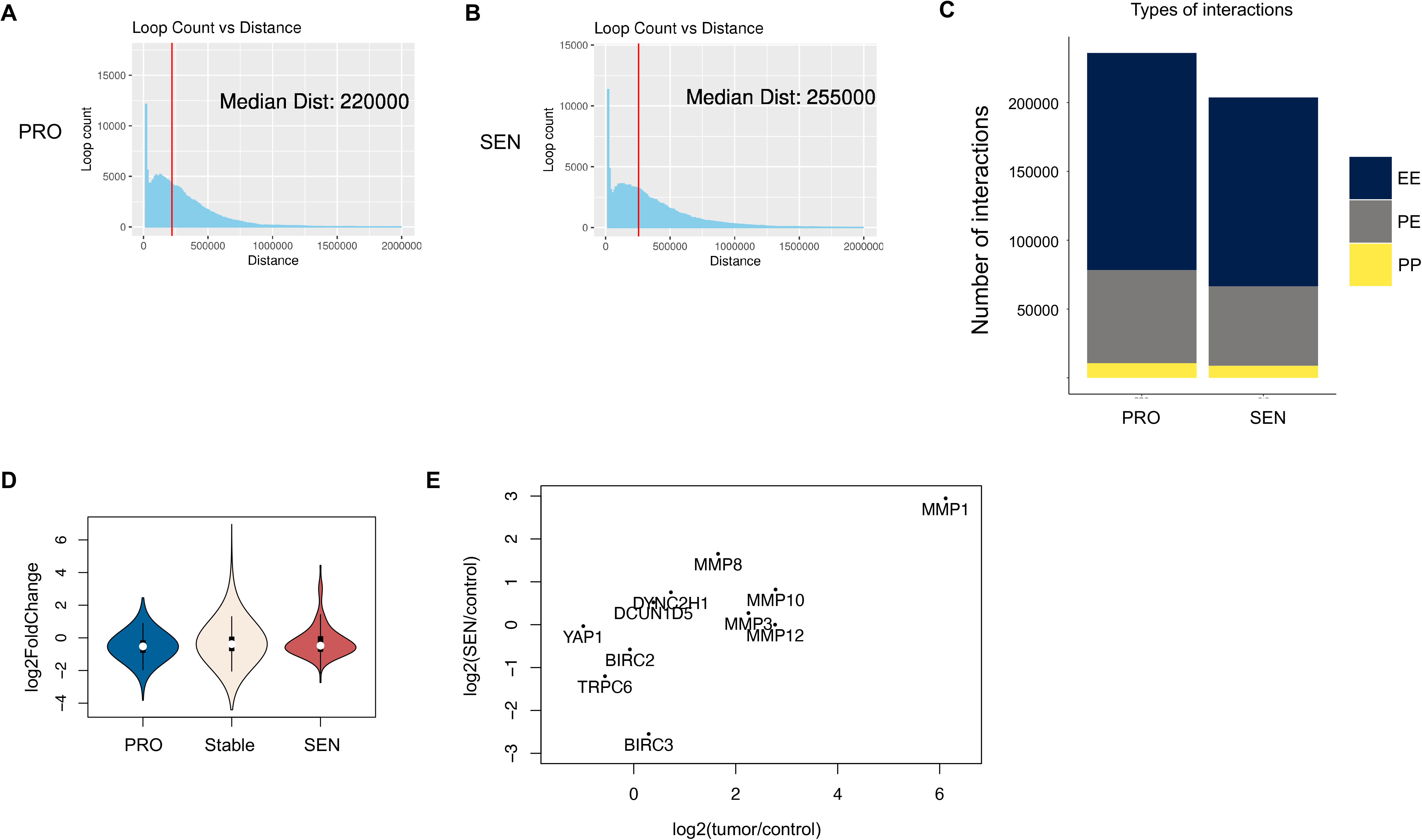
Enhancer connectome in proliferating and senescent cells. A. Distribution of looping distance in proliferating (PRO) cells. B. Distribution of looping distance in senescent (SEN) cells. C. Distribution of different types of enhancer interactions. D. CTCF binding in proliferating-specific, stable, and senescence-specific enhancer communities. E. Matrix metalloproteinase and other genes in a SEN-specific enhancer community exhibit a linear correlation in gene expression increase in SEN and breast cancer. PRO, proliferating cells; SEN, senescent cells.

**Supplemental Figure 3.**
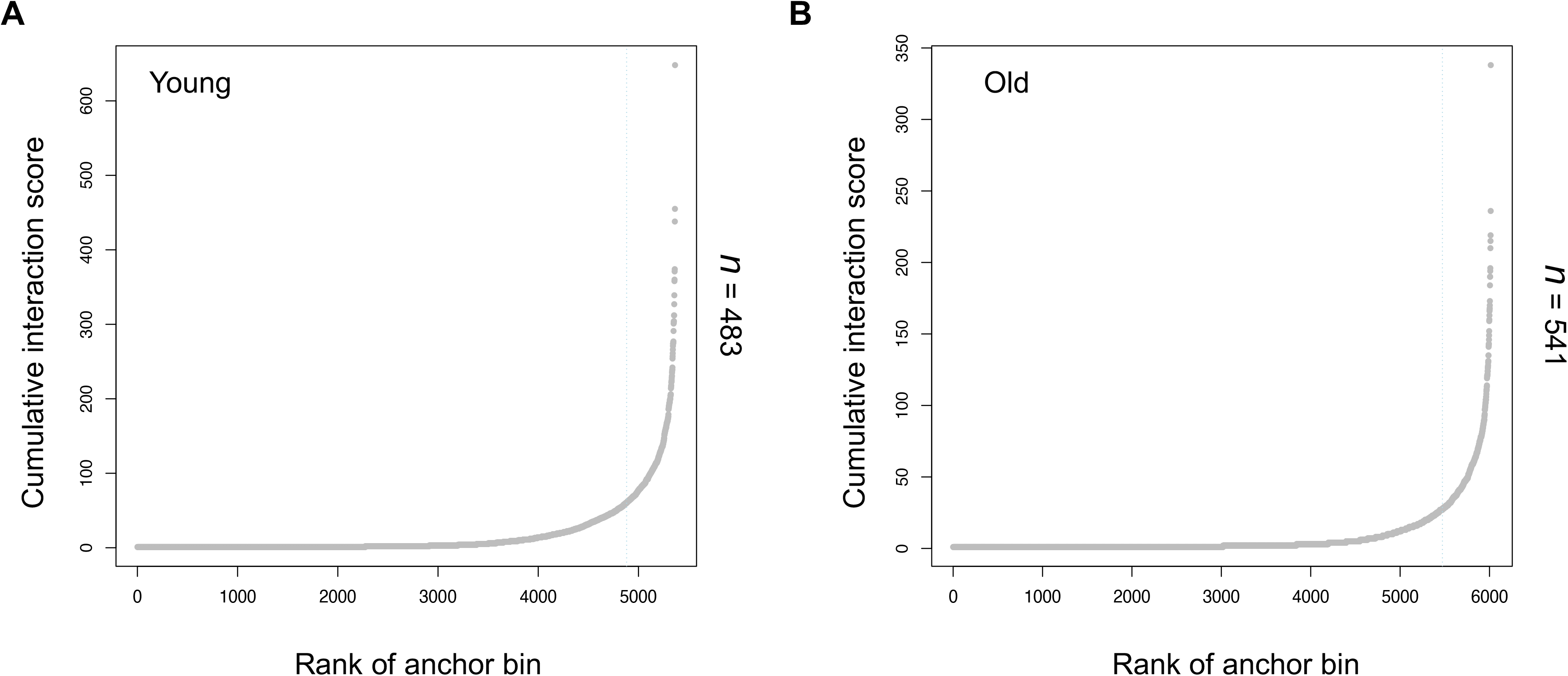
Enhancer connectome in young and old mice. A. Hyper-connected enhancer communities in hepatocytes isolated from young mice. B. Hyper-connected enhancer communities in hepatocytes isolated from old mice.

**Supplemental Table 1.** H3K27ac HiChIP processing.

| Sample | PRO_rep1 | PRO_rep2 | SEN_rep1 | SEN_rep2 |
| --- | --- | --- | --- | --- |
| Total pairs | 544,343,426 | 511,403,209 | 456,557,641 | 606,028,982 |
| Unique alignments | 396,140,052 | 364,481,762 | 325,329,957 | 434,740,543 |
| Valid interactions | 335,976,267 | 310,688,049 | 289,303,228 | 386,696,676 |
| Cis interactions | 245,123,000 | 213,093,572 | 202,516,717 | 272,715,066 |

## Methods

### Cell culture and treatment

Primary IMR90 fibroblasts were cultured in DMEM supplemented with 10% fetal bovine serum (FBS), 100 units/mL penicillin, and 100 μg/mL streptomycin (Invitrogen). Cells were routinely tested for mycoplasma. The cells were cultured under physiological oxygen (3%), and were used within population doubling of 35. For HRasV12-induced senescence, retrovirus from retroviral vector encoding HRasV12 was used to infect cells. Following hygromycin selection, cells were cultured for 7 or more days.

### Mice experiments

Male, 3-month-old C57BL/6 mice were purchased from Charles River Laboratories (Wilmington, MA) and 23-month-old male mice from the US National Institute on Aging stock located in Charles River Laboratories were used in this experiment. Mice were maintained at Sanford Burnham Prebys vivarium for 2 months prior to experiment to allow mice to acclimate. Mice were maintained under a specific pathogen-free environment and kept under standard conditions with a 12-h light/dark cycle and fed ad libitum on a regular diet. All animal experiments were carried out under procedural guidelines of Institutional Animal Care and use Committee (IACUC).

### Hepatocyte isolation

Mice were anesthetized, and the livers were perfused with perfusion media through the portal vein and then digested with digestion media with collagenase I. Digested tissue was suspended with DMEM media, and the suspension was passed through a 100 μm nylon mesh filter (BD Falcon). Suspended tissue was centrifuged at 50 g for 3 min to pellet hepatocytes. Red blood cells were removed. Cells were washed with PBS, and then used for downstream experiments.

### ATAC-seq experiments

Omni-ATAC libraries were prepared following the protocol previously described. Briefly, 50,000 cells were resuspended in the following buffers after sequential centrifugation, 1 ml of resuspension buffer (RSB), 50 μl of RSB containing 0.1% Tween-20, and 50 ul of transposition mix (25 μl of 2X TD buffer, 2.5 μl of TDE1, 16.5 ul PBS, 0.5 μl of 1% digitonin, 0.5 UL 10% Tween-20, and 5 μl of nuclease-free water). The transposition reaction was then incubated at 37 °C for 30 min in a Thermo Mixer with agitation at 1000 rpm. The reaction was cleaned up using a QIAGEN MinElute Reaction Cleanup kit. Transposed fragments were amplified and purified as described previously. Libraries were sequenced in paired-end mode on a NextSeq 550 instrument.

### PRO-seq experiments

qPRO-seq was performed as previously described^51^. Briefly, cells were permeabilized using permeabilization buffer and checked with Trypan blue. Permeabilized cells were washed using cell wash buffer and stored in cold freeze buffer at −80°C until use. The permeabilized cells were then resuspended in 2X run-on master mix containing biotin-11-CTP, ATP, GTP and UTP for 5 minutes at 37C. RNA was extracted using Norgen total RNA purification kit (Norgen, 37500), and then subjected to 3’ RNA adaptor ligation and streptavidin bead binding. The 5’ ends of the RNA were repaired, decapped, and ligated to 5’ adapter and subjected to TRIzol elution. The RNA was then reverse transcribed, and the number of PCR cycles were determined based on qPCR. The final libraries were then amplified with appropriate index primers. The qPRO-seq libraries were pooled and sequenced in pair-end mode on the NextSeq 550 platform.

### HiChIP experiments

The HiChIP experiments were carried out following the protocol previously described^18^, using the H3K27ac antibody (Abcam, ab4729). Briefly, 10 million cells were crosslinked using 1% formaldehyde for 10 minutes, and then sonicated for 6 minutes. After the immunoprecipitation step, DNA were purified and then quantified using Qubit. The amount of Tn5 used and the number of PCR cycles were determined as previously described based on Qubit quantification^18^. After PCR amplification, size selection was performed using Ampure beads. Libraries were sequenced in pair-end mode on a NextSeq 550 instrument.

### ATAC-seq data processing

Quality control checks were conducted on raw reads using FastQC, and adaptor sequences were trimmed by fastp. Then, reads were aligned against hg19 using bowtie2 with parameters “--very-sensitive -X 2000”. Reads mapped to mitochondrial DNA were removed. Properly paired reads with MAPQ more than 30 were kept and duplicates were removed using Picard and samtools. Technical replicates were merged, and samples were downsampled to similar coverage. Reads were shifted. Peaks were called using macs2. Reproducible peaks between biological replicates were used for downstream analyses.

### Cut&Run data processing

Cut&Run data processing followed the published protocol^35, 52^. Quality control checks were conducted on raw reads using FastQC, and adaptor sequences were trimmed by Trimmomatic. Then, paired reads were aligned against hg19 using bowtie2 with parameters “--dovetail”. Properly paired reads with MAPQ more than 2 were kept using samtools. Duplicates were removed using Picard. Peaks were called using macs2.

### qPRO-seq data processing

qPRO-seq data were processed following previously published protocol^51^. Briefly, quality control checks were conducted on raw reads using FastQC, and adaptor sequences were trimmed using fastp. rDNA reads were excluded from the downstream analyses. Then, paired reads were aligned against hg19 using bowtie2. Duplicates were removed using UMI-tools. Bigwig files were generated based on strand information.

### HiChIP data processing

Paired-end reads were aligned to hg19 genome using HiC-Pro with default parameters for MboI restriction fragments^53^. Duplicates were removed using Picard. Sequencing depth and alignment information for each library were summarized in Supplemental Table 1. Significant interactions were called using FitHiChIP at the resolution of 5kb bins^54^.

### Hyper-connected community analysis

The hyper-connected community were called following the previously reported procedure^14, 15^. Each vertex was an anchor, and each edge was a significant interaction. The hyper-connected communities were called by the cluster_louvain function in the R packge igraph with default parameters. The connectivity of these communities was ranked and the cutoff was determine as the elbow of the curve.

### Motif analysis

Enriched transcription factor binding motifs were identified within proliferating- or senescence-specific hyper-connected communities using HOMER v4.6 findMotifsGenome.pl with -size given. Results were mined from the known motif (supervised) assessment. Scanning windows were obtained by taking the anchor points of every community in a given condition (e.g., senescence) with no anchor point overlaps to any cliques in the opposite condition (e.g., proliferating), then crossing these to the set of all ATAC-seq open chromatin regions from any condition. The background was set to be scanning windows from the opposite condition.

### MafK binding analysis

Senescence-specific MafK CUT&RUN peaks were identified using bedtools intersect - wa -v, and these were intersected with senescence-specific anchor points using bedtools intersect -wa -u to produce a list of senescence-specific loops/peaks. Each differentially expressed gene was then assessed for the number of connected SEN-specific loops/peaks using its enhancer community. PRO-seq log2(SEN/PRO) gene upregulation was then plotted in box-and-whisker, separating "high" (> 20 SEN-specific loops/peaks) from "moderate" (<= 20 SEN-specific loops/peaks) MafK-bound genes, with the background formed by differentially expressed genes which do not have specific loops/peaks. Pro-specific MafK loops/peaks were assessed in reverse of description above. Statistical significance was assessed using the R package coin’s independence_test() function (permutation testing).

### GWAS analysis

Fold enrichment for GWAS categories was obtained by overlapping all SNPs from the NHGRI-EBI GWAS catalog to PRO-specific, SEN-specific, or shared anchor points, and then calculating (e.g., for SEN-specificity):

(# SEN-specific anchor points with SNPs / # total SEN-specific anchor points) / (# shared anchor points with SNPs / # total shared anchor points)

GWAS terms with an SEN-specific fold enrichment > 1.5 and 10 or more SEN anchor points with SNPs were plotted for fold enrichment (size) and # anchor points with SNPs (heat), sorting in ascending order by the difference in fold-enrichment SEN-Proliferating.

### TCGA analysis

Breast cancer data were downloaded from TCGA through the GDC data portal and normalized TPM measurements of gene expression were collected for 1,427 primary tumors and 113 solid tissue control samples. Breast cancer was chosen for study because a selection of upregulated SASP genes in enhancer communities which grow in SEN and include novel MafK binding events tend to be mutated most commonly in cancers of this tissue (OncoPrint tool)^55^. Tumor and control values were averaged over all samples and a score log2(tumor/control) was assessed for each gene. Enhancer communities were scored based on the correlation of TCGA tumor upregulation and SEN upregulation (RNA-seq from GEO GSE70668^56^), considering targeted genes which have a net gain of > 5 loops in SEN; enhancer communities were deemed to be "cancer-correlated" if the Spearman rank correlation was > 0.6 or "cancer-anticorrelated" if it was < -0.6. Genes targeted by these enhancer communities were then mined for Gene Ontology term enrichment (DAVID, FDR < 0.1). As a control for new looping, enhancer communities were also assessed using all genes and examining the correlation, with subsequent GO term enrichment analysis. Cancer-correlated or - anticorrelated communities were evaluated for motif enrichment in the following way: anchor points from cancer-correlated enhancer communities were filtered to remove any which overlap with gene bodies or promoters (1kb upstream regions from RefSeq TSSs). The resulting distal intergenic anchor points were then crossed with ATAC-seq open chromatin regions, yielding likely enhancers. These were scanned using HOMER v4.6 findMotifsGenome.pl with a fixed size 300bp window and repeats masking (-size 300 -mask) against a background of correlated communities from the opposite type: senescence correlated vs proliferating correlated, proliferating correlated vs senescence correlated, senescence anti-correlated vs proliferating anti-correlated, proliferating anti-correlated vs OIS anti-correlated.

## Data Availability

Data generated in this study have been uploaded to NCBI GEO under series accession number GSE232942.

## Acknowledgements

We thank Y. Lan for help with the HiChIP pipeline. L.W. is supported by NIH grant K99AG065500. P.D.A. and S.L.B are supported by NIH grant P01AG031862.

## Author contributions

L.W. and S.L.B. conceived the project. L.W. performed most of the experiments and analyses. L.W. and C.X. performed cell-culture and knock-down experiments. L.W. and C.Z. assessed senescence phenotypes. A.H. and P.D.A contributed to the mouse experiments. L.W., G.D., X.L., W.W., G.V. contributed to the analyses. L.W. and S.L.B. wrote the manuscript with input from other authors. All authors discussed the results and reviewed the manuscript.

